# Differential gene expression analysis of spatial transcriptomic experiments using spatial mixed models

**DOI:** 10.1101/2024.01.20.576348

**Authors:** Oscar E. Ospina, Alex C. Soupir, Roberto Manjarres-Betancur, Guillermo Gonzalez-Calderon, Xiaoqing Yu, Brooke L. Fridley

**Affiliations:** Department of Biostatistics & Bioinformatics, H. Lee Moffitt Cancer Center and Research Institute, Tampa, FL, USA; Biostatistics and Bioinformatics Shared Resource, Moffitt Cancer Center, Tampa, FL, USA; Division of Health Services & Outcomes Research, Children’s Mercy, Kansas City, MO, USA

## Abstract

Spatial transcriptomics (ST) assays represent a revolution in the way the architecture of tissues is studied, by allowing for the exploration of cells in their spatial context. A common element in the analysis is the delineation of tissue domains or “niches” followed by the detection of differentially expressed genes to infer the biological identity of the tissue domains or cell types. However, many studies approach differential expression analysis by using statistical approaches often applied in the analysis of non-spatial scRNA data (e.g., two-sample t-tests, Wilcoxon rank sum test), hence neglecting the spatial dependency observed in ST data. In this study, we show that applying linear mixed models with spatial correlation structures using spatial random effects effectively accounts for the spatial autocorrelation and reduces inflation of type-I error rate that is observed in non-spatial based differential expression testing. We also show that spatial linear models with an exponential correlation structure provide a better fit to the ST data as compared to non-spatial models, particularly for spatially resolved technologies that quantify expression at finer scales (i.e., single-cell resolution).

## Introduction

The ability to measure gene expression within a spatial context, which is referred to as spatial transcriptomics (ST), includes a wide range of technologies, including assays based on the well-stablished in-situ fluorescent hybridization (FISH)^1–3^, and groundbreaking in-situ spatial barcoding^3–8^. Current ST techniques have the capacity for large multiplexing (i.e., hundreds to thousands of genes assayed in the same tissue) and the generation of an additional data modality representing the spatial position of the measured gene expression. The spatial information from ST experiments has allowed researchers to address questions about the tissue architecture of organs and diseases^3,9–11^. Of special importance has been the use of ST to assess tissue heterogeneity in many cancerous tissues^6,12–21^, as well as infected tissues^22^. Spatial transcriptomics has also enabled a better understanding of cell-to-cell communication^23–25^ and the identification of potential druggable targets^18,26,27^.

One common step in ST analysis is the identification of genes that differentiate tissue domains within a sample (i.e., differentially expressed genes among tissue niches)^28–30^. Because of the similarity in statistical data properties between scRNA-seq and ST, many studies complete the identification of ST gene expression within domains in an analogous fashion as it is carried out among scRNA-seq cell clusters or cell populations. Once tissue niches have been identified in the ST samples, researchers often proceed with non-parametric tests such as the Wilcoxon rank test^31–33^ to identify differentially expressed (DE) genes among the niches. Although this approach may be appropriate for cases where transcriptomic differences between the compared domains are large (e.g., tumor vs. stroma), it does not account for the spatial dependency, which results in gene expression of neighboring sampling units (e.g., cell or spots) to be more similar than distant sampling units^34^. Because the spatial dependency in ST data is a driving factor of the gene expression patterns observed in tissues^35,36^, more sophisticated statistical methods could be used to account for the spatial dependency between sampling units^37–39^. Common approaches in many novel methods include identifying genes with spatial patterns, such as gene expression “hot spots”, or testing for genes showing high expression on each tissue domain (i.e., cluster) detected in a sample^35,38–44^. Benchmarking to compare the performance of these approaches has also been done^45^, which is crucial to aid in method selection. However, despite the wide availability of methods to detect spatially variable genes, less effort has been directed to quantify the impact of disregarding spatial dependency in ST data analysis.

Quantifying the impact of non-spatial approaches for the detection of differentially expressed genes is an important endeavor, given that failure to account for the spatial autocorrelation in ST experiments may result in inflation of the type I error rate^40,46–48^. An increased type I error rate leads to more genes erroneously being identified as differentially expressed due to inaccuracy in the p-values (i.e., p-values too small). The impact of inflated type I error rates is increased due to unreliable estimation of gene expression variation, as the variation estimates do not consider the spatial correlation among the neighboring and distant sampling units.

The use of linear mixed models offers a simple alternative for DE analysis in ST data. In bulk RNA-seq analysis, robust and well-established pipelines apply linear model fitting to test for differences in expression between two or more categories^49,50^. However, their application to ST requires additional considerations, given the spatial nature of this modality. One such consideration, which takes advantage of the flexibility of linear mixed models, is the incorporation of spatial covariance structures and variogram analysis^51,52^. To implement this approach as an alternative for the analysis of ST data we introduce STdiff, a method for differential gene expression analysis among groups of sampling units in ST experiments. The method tests for genes with significantly higher (or lower) expression in one group (e.g., cluster, tissue niche) with respect to the others by fitting linear mixed models that explicitly account for the random spatial effects via spatial covariance structures. To demonstrate the utility of STdiff, we tested for DE genes in publicly available ST data sets generated with 10X Genomics’ Visium platform and Nanostring’s GeoMx and Spatial Molecular Imager (SMI) platforms. We fitted corresponding non-spatial and spatial models to assess the impact of accounting for the spatial autocorrelation on the downstream DE analysis results.

## Results

### Comparison of non-spatial and spatial models

Models with or without spatial covariance structures were fitted for each gene to determine the most suitable alternative for capturing the expression differences among tissue domains. The tissue domain or cell type annotations for each region of interest (ROI), spot, or cell were obtained from the studies that generated the data sets (**Table 1**; **Supp. Table 1**). The studies generated the annotations using histopathology methods (Visium and GeoMx data sets) and cell phenotyping (CosMx data sets). Assessment of the models using the Akaike Information Criterion (AIC) showed that spatial models with an exponential covariance structure provided a more accurate fit to Visium and SMI data than non-spatial models (**Fig. 1**). Among the four Visium samples, between 28–41% of the tests (i.e., gene expression in domain A vs gene expression in other domains) showed a better fit to the data when using a spatial model (i.e., lower AIC) compared to a non-spatial model. For the SMI datasets, the percentage of tests favoring the spatial models varied from 32–67%. In contrast, for the analysis of the GeoMx data sets, no more than 16% of the spatial models were favored over the non-spatial models (**Fig. 1**). When considering only genes with high expression in the samples (above the median expression), the proportion of favored spatial models increased to 48–66% in Visium studies and 51–93% in SMI studies (**Fig. 1**).

**Figure 1.**
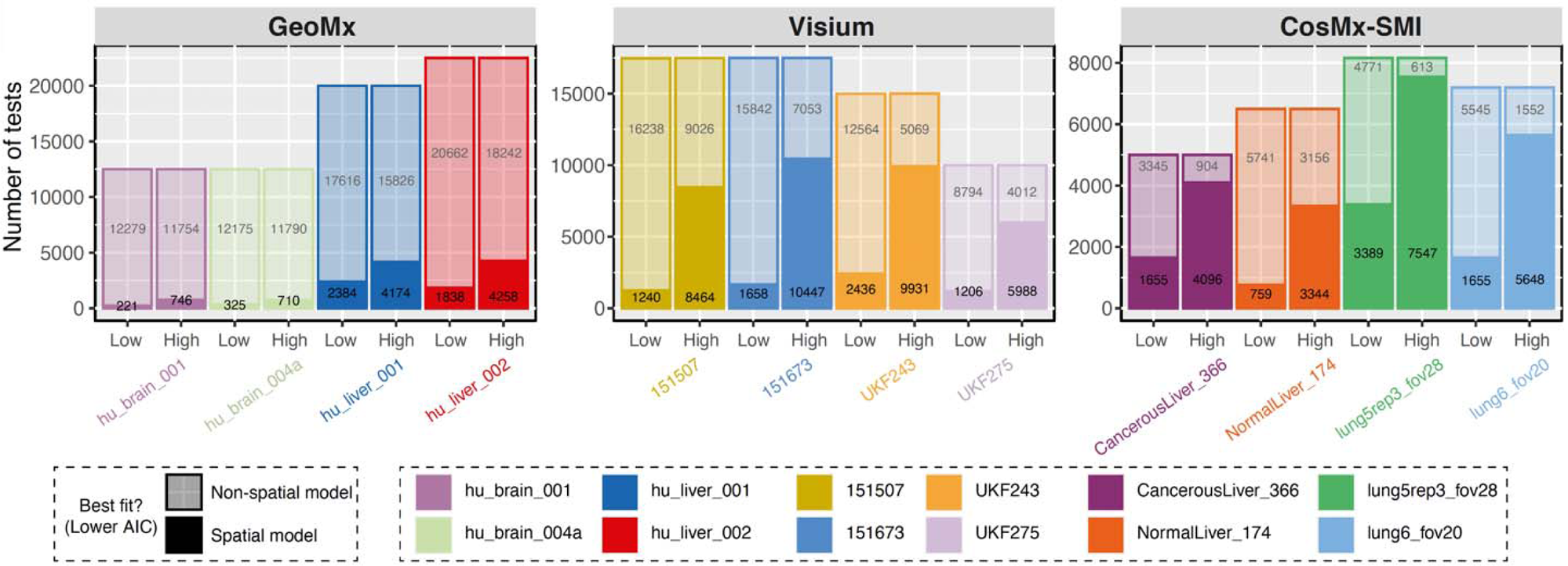
The results of model comparison between non-spatial models and spatial models with exponential covariance structure using AIC. For each gene x cluster test, the models with the lowest AIC were deemed to be a better fit to the data (solid color: spatial model with lower AIC, translucid color: non-spatial model with lower AIC). The tests were separated according to the average gene expression across all ROIs/spots/cells in the tissue sample (high vs low expression based on the median gene expression as threshold)

**Table 1.**
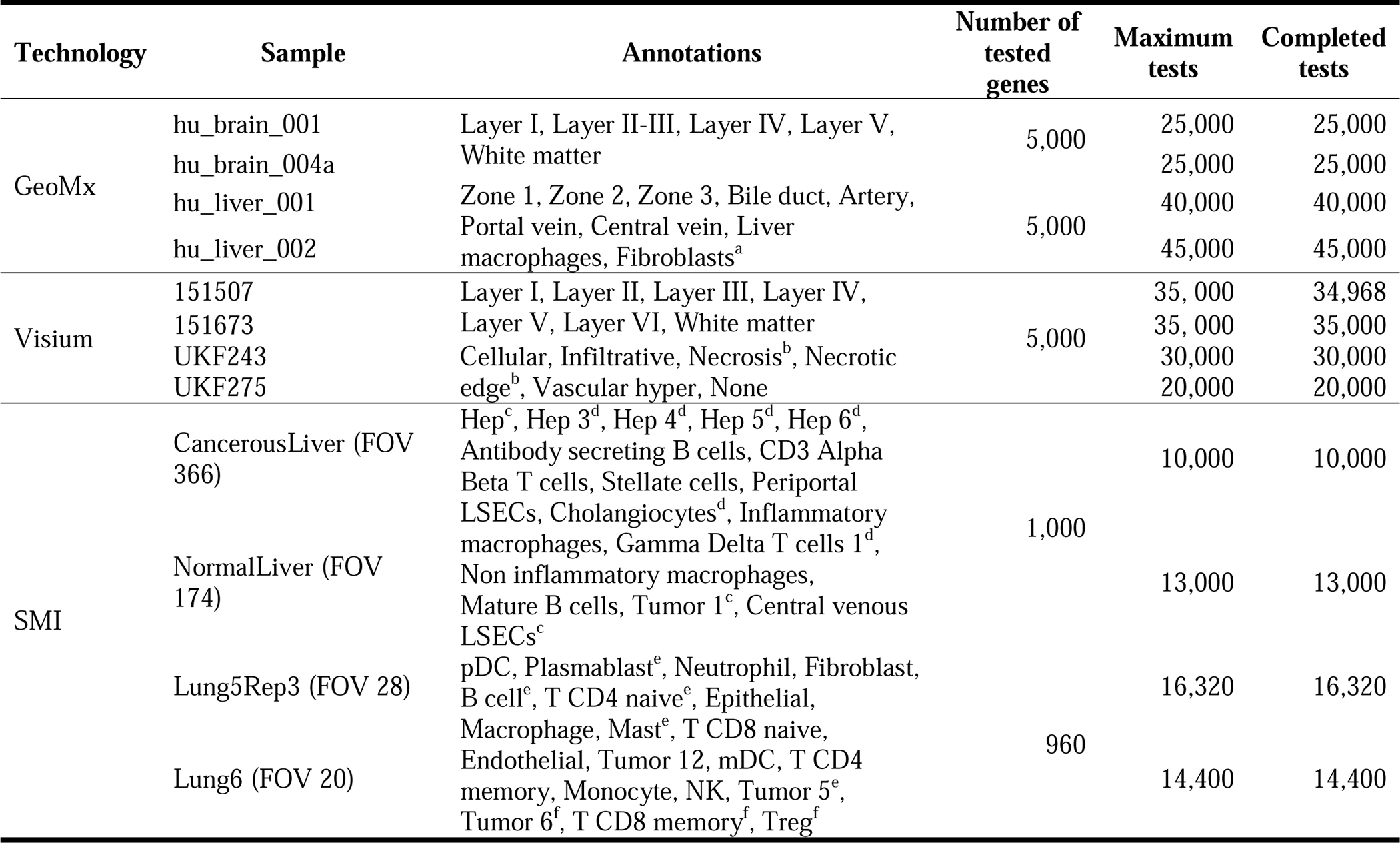
Summary of spatial transcriptomics samples used in the differential expression tests. The biological annotations present in each sample are also shown. The maximum number of tests performed corresponds to the combination of genes times the number of annotations. The completed tests column indicates the number of tests that reached convergence during the REML optimization in spaMM (using the exponential covariance structure). ^a^ Not present in sample hu_liver_00. ^b^ Not present in sample UKF275. ^c^ Not present in sample NormalLiver (FOV 174). ^d^ Not present in sample CancerousLiver (FOV 366). ^e^ Not present in sample Lung5Rep3 (FOV 28). ^f^ Not present in sample Lung6 (FOV 20)

### Control of type I error by spatial models

The DE p-values tended to be smaller in the non-spatial models compared to the spatial models, possibly due to an increase in the type-I error inflation. However, these patterns were dissimilar among the ST technologies (**Fig. 2**). In the case of Visium experiments, 65–71% of the p-values were larger in the spatial models compared to the non-spatial models. In SMI, 60–66% of the p-values from the spatial models were larger than those from the non-spatial models. In the GeoMx experiments, the p-values from the spatial models were larger in 40–54% of the tests compared to the non-spatial models. These modeling results suggest a potential slight inflation in the type I error rate for the non-spatial models, whereby p-values generated by non-spatial models are too small likely due to inaccurate estimation of the variance in test statistic. In other words, the variance estimation for the non-spatial models is too small, resulting in a larger test statistic and artificially smaller p-value.

**Figure 2.**
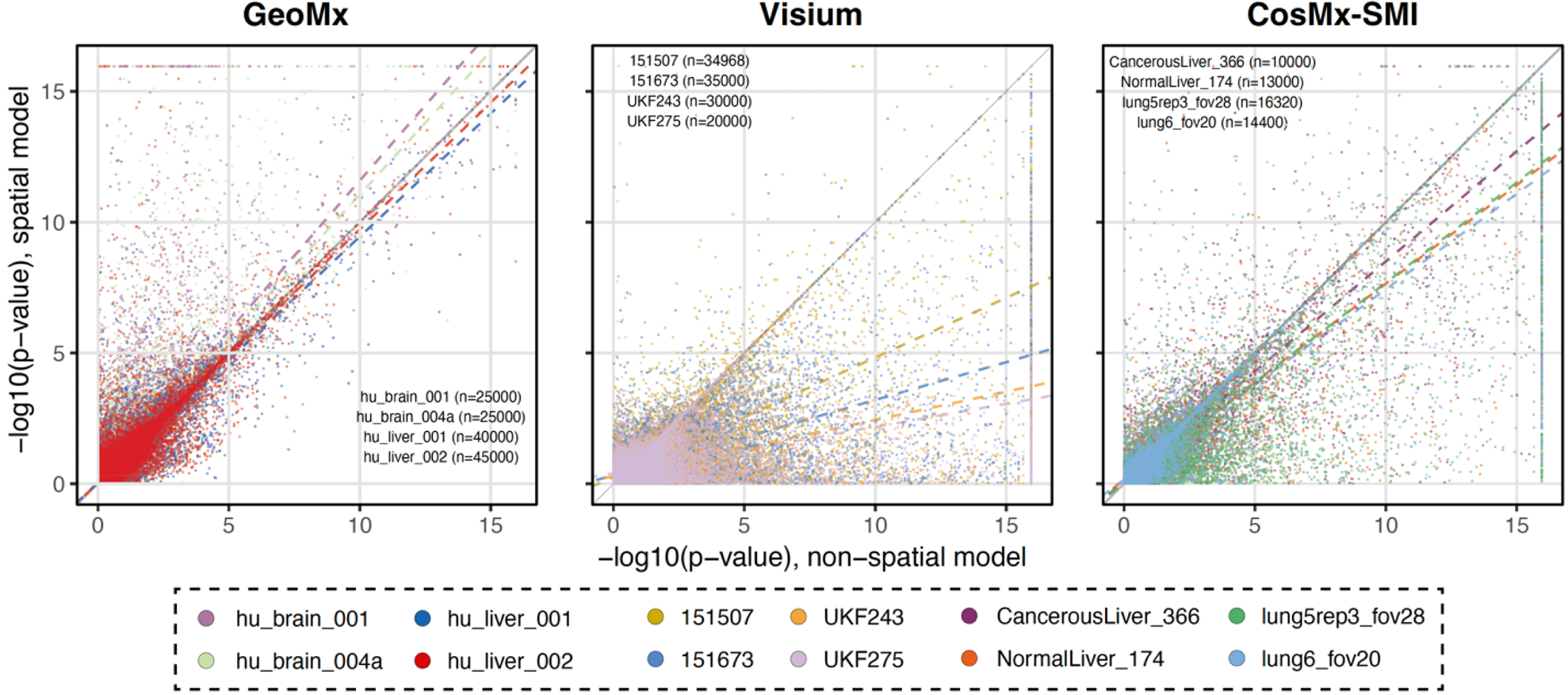
Comparison between non-spatial and spatial (exponential model) differential expression tests. Each point corresponds to the −log10(p-value) resulting from a (non-spatial or spatial) linear model fit between the expression of a gene and a binary variable indicating whether a ROI/spot/cell belongs to a biological annotation. The p-values indicate if the gene is differentially expressed (model coefficient different to zero) for a specific biological annotation with respect to the rest of the ROIs/spots/cells. The solid line indicates a 1:1 correspondence (i.e., non-spatial and spatial models yield the same p-values). The colored dashed lines indicate the linear trend of the p-values for each sample. If a colored line lies below the solid line, p-values from the non-spatial model tend to be larger than those from the spatial model.

## Discussion

Researchers often aim to detect differences in gene expression between cells or tissue niches, with many methods available for non-spatially informed assays such as single-cell or “bulk” RNAseq^49,50,53,54^. Although spatial statistics methods have existed in the literature for several decades^51^, only recently have spatial statistics been applied to the detection of spatially variable genes in biological tissues assayed with ST^35,38–44^. In this study, we have shown that the detection of differentially expressed genes in ST data benefits from statistical models that consider spatial autocorrelation. This leads to a more accurate estimate of the variance and thus produces more stable estimates of p-values. In other words, the spatial models account for the non-independence in the cells/spots, which is not addressed by traditional non-spatial linear models (i.e., two sample t-tests assuming independence between observations). Failure to consider this dependency between observations may cause the tests to underestimate the variance of the test statistic resulting in overly small p-values. Our results highlight the importance of considering the spatial dependency present in spatial-resolved transcriptomics data, which is often neglected in many studies in conducting differential expression analyses.

Our results comparing the models with and without a spatial correlation structure indicated that for densely sampled ST data (e.g., Visium, SMI) spatial models present a better model fit. For non-densely sampled experiments (e.g., GeoMx using ROIs), there was a slight tendency for non-spatial models to fit the data better when compared to spatial models, probably due to less spatial correlation among ROIs that are often sampled distant from one another. In light of this finding, the use of non-spatial models, such as two-sample t-tests, may be appropriate to study differential gene expression in studies using GeoMx where the ROIs are more spatially distant. Nonetheless, the correlation among ROIs within a single slide and the technical variation among slides in the same study could be taken into account when testing for differentially expressed genes by accounting for the spatial autocorrelation^55^. Our results also indicate that for Visium and SMI, the spatial models performed better compared to non-spatial models in cases where the differential expression test involved a highly expressed gene. Genes with low expression are likely to show excessive zeroes (a characteristic of ST data^56,57^), and hence, fitting spatial mixed models may become challenging. Novel application of Bayesian methods to detect spatially variable genes appears to be robust to excessive zeroes in ST data^57,58^.

Our results were indicative that p-values obtained from the spatial model constituted a more biologically informative ranking metric for gene set enrichment analysis (GSEA). Using Benjamini-Hochberg (FDR) adjusted p-values from the non-spatial and spatial models as ranking metrics we performed GSEA for the Hallmark gene sets with the R package fgsea^59^ ^60^. The GSEA was conducted individually for each histopathology-defined domain in the glioblastoma Visium data set^61^. We observed that across all the significantly enriched Hallmark gene sets, the results were more significant using the p-values from the spatial models as compared to the non-spatial models, with the exceptions of oxidative phosphorylation in the necrosis niche and KRAS signaling downregulation in the necrotic edge niche (**Fig. 3**). A lower score of the KRAS signaling is expected in the necrotic edge, under the assumption that the tumor cells in this niche are not actively proliferating^62^. Although the GSEA was conducted on a single Visium sample (UKF243), and comprehensive testing is required to evaluate the information p-values can provide for pre-ranked GSEA, our analysis suggests that p-values derived from spatial models can be more appropriate for gene set enrichment analysis when using ST data.

**Figure 3.**
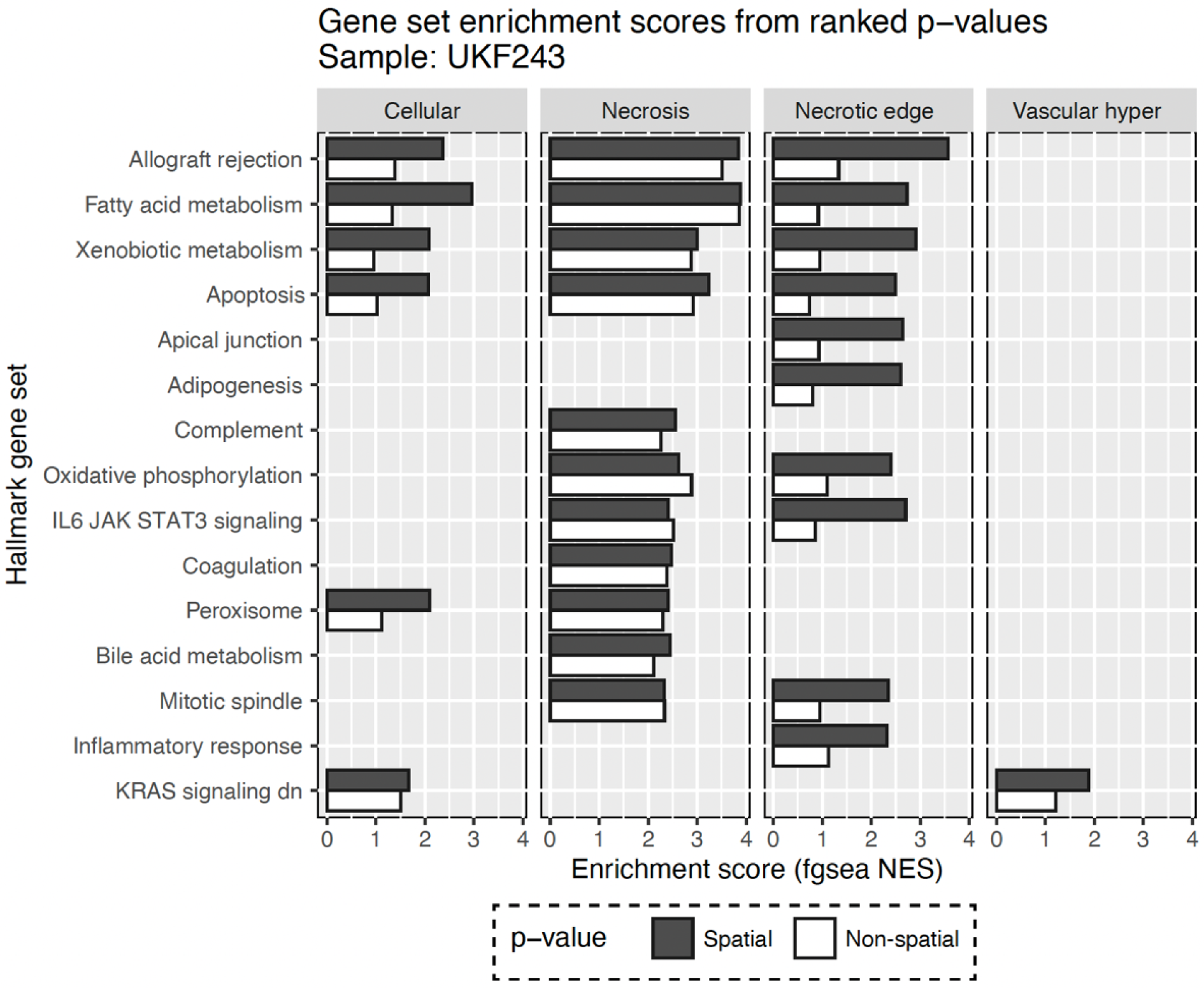
Scores resulting from the GSEA analysis calculated using the fgsea package. Genes were ranked using the p-values obtained by the non-spatial (white) or non-spatial models (dark grey). The spatial niches (cellular, necrosis, necrotic edge, vascular_hyper) were generated via histopathology examination by a previous study^61^. The gene sets depicted here represent Hallmark gene sets showing significant enrichment (adjusted p-value <0.05). The “vascular_hyper” niche refers to tumor tissue with high vascularization.

Testing for differential gene expression is time-consuming for modern single-cell or spatial applications, as hundreds to thousands of individual tests are performed (i.e., each combination of gene expression in domain A vs gene expression in other domains). In addition, each test often includes hundreds to thousands of cells or spots. When applying spatial models for differential expression, the advantages of accurate estimation come at the cost of longer computation times compared to the non-spatial models (**Fig. 4**). Previously, we performed these models using the long-supported R package *nlme*. However, the estimation of parameters was exceedingly time consuming (data not shown). Hence, we switched to using the R package *spaMM* to fit the statistical models. Using a High-Performance Computing environment (HPC) it can take anywhere from a few seconds to up to more than 2 hours to test for one gene between two tissue domains in Visium- or SMI-generated data. We ran our test using a single core per each test and 8GB of memory, which are resources not typically available in conventional laptop computers. After considering these results, we opted to implement differential gene expression analysis using *spaMM* (as opposed to *nlme*) in our R package for spatial transcriptomics analysis *spatialGE*^63^, and we have named this approach STdiff. In the *spatialGE* R package, we made efforts to parallelize the analyses, but such efforts alone are not enough to achieve feasible computing times on personal computers and require the use of an HPC environment.

**Figure 4.**
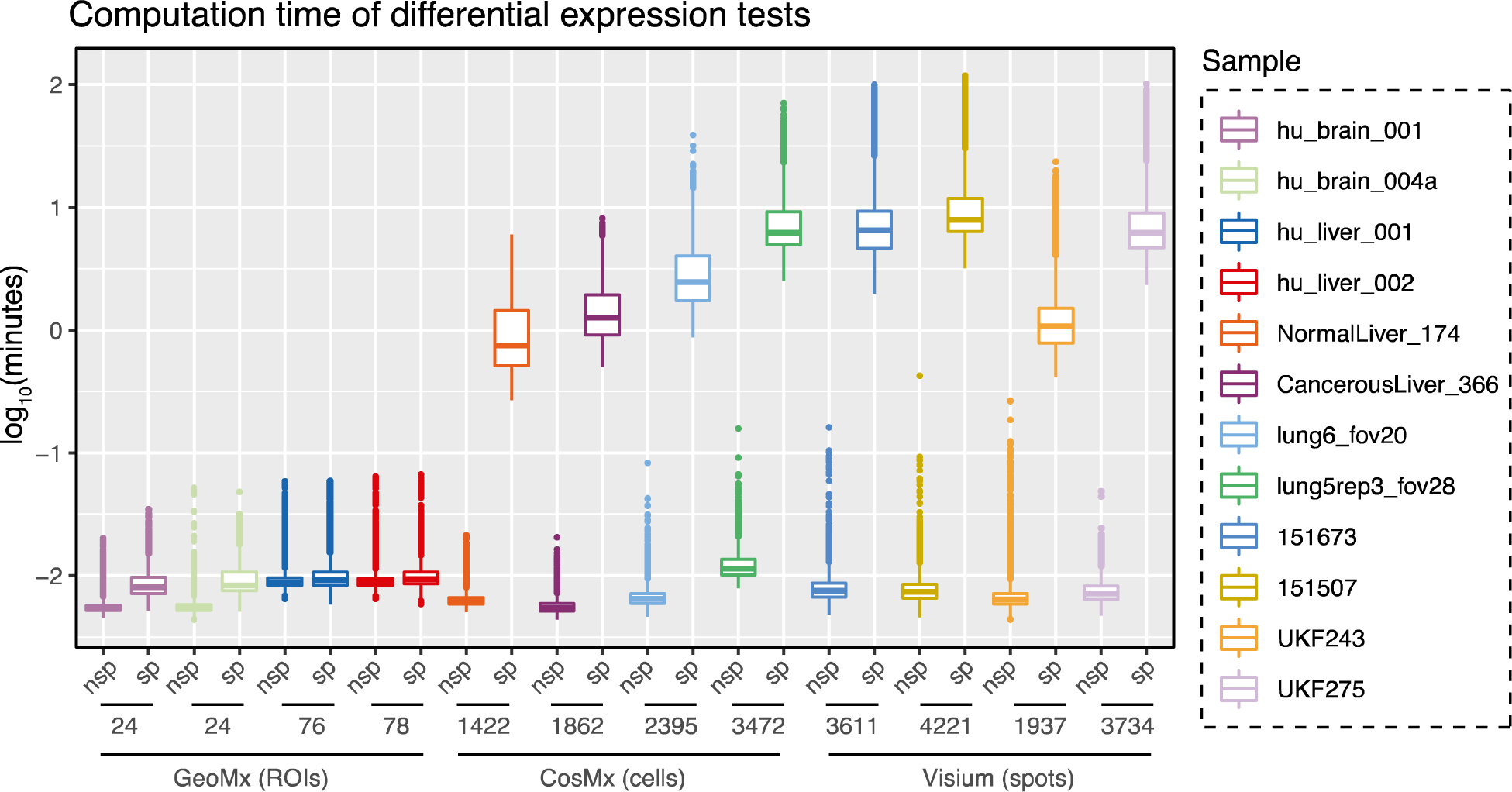
Time of execution in log_10_(minutes) for the non-spatial (nsp) and spatial (sp) test conducted. Each dot represents a gene x cluster test. The number of ROIs, spots, or cells is shown on the x axis.

In summary, consideration of spatial dependency is needed when conducting differential expression analysis in densely sampled spatially resolved transcriptomic experiments. In this study, we demonstrate that applying mixed models with spatial correlation structure effectively accounts for the correlation between spots or cells, thereby controlling for the inflated type I error rates observed in non-spatial models. Specifically, we show that spatial models with an exponential correlation structure provide a better fit to ST data than non-spatial models.

## Material and Methods

### Spatial transcriptomic data sets

Spatial transcriptomics technologies are diverse, ranging in cellular and molecular resolution. Hence, we tested the utility of spatial linear mixed models for differential gene expression analysis using a series of data sets that reflected the spectrum of cellular and molecular resolution in ST technologies. We obtained publicly available ST data from spatial-barcoding technologies, including 10X Genomics’ Visium and NanoString’s GeoMx platforms, as well as the imaging technology produced from NanoString’s CosMx Spatial Molecular Imager (SMI). The Visium data sets were generated by previous studies of the brain motor cortex^64^ and glioblastoma^61^. The GeoMx and SMI data sets were obtained from NanoString’s Spatial Organ Atlas repository^65^. For each technology, we selected two tissue types with two samples for each tissue type (i.e., a total of 4 samples for each technology). More details of the selected samples and their access links are provided in the supplemental materials (**Supp. Table 1**). Using these data sets, we tested the utility of spatial models to detect DE genes. For this reason, a requisite for sample selection was that it contained biologically meaningful annotations (i.e., tissue domains, niches, or clusters) for each ROI/spot/cell. Preparation of expression and annotation data was carried out using the R statistical programming software version 4.1^66^. Data was normalized using library size normalization and log-transformation in the package spatialGE^67^.

### Model

In differential gene expression analysis, the goal is to identify genes for which the average expression in a group is significantly higher or lower than that in other groups. In the context of ST, the sampling units (cells, spots, ROIs) are grouped using either a clustering method or prior knowledge of the tissue (e.g., tissue domains or niches). Hence, the objective remains the same: To detect genes with significantly higher or lower expression in one group of cells, spots, or ROIs (i.e., spots or cells in a domain or tissue niche) compared to ROIs/spots/cells in another tissue domain or outside of the tissue domain of interest.

For the non-spatial case of our DE analysis proposal, the expression of a given gene (*y_s_*) at a given sample unit location (*s*) can be modeled as:

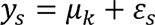

where *µ_k_* is the mean expression of the gene in cluster *k*, and *ε_s_* is the random error at location *s*, with *ε_s_* ∼*N*(0, σ^2^). In order to extend this model to the spatial case, we add the effect of the spatial dependency as part of the random effects (*U_s_*) term to account for the correlation among neighboring sampling units as:

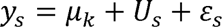

where *U_s_* is defined as *U_s_* ∼*MNV*(0, *V*(*θ*, *d*)), where d represents the distance between two ROIs/spots/cells. The spatial dependency can be defined by several types of covariance structures. In this study, we have tested the use of the commonly used exponential covariance structure which is a special case of the Matérn covariance structure, 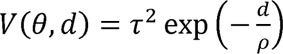.

### Application of models on spatial transcriptomic data sets

The application of spatial models to densely sampled tissues can be computationally intensive, particularly as the number of ROIs/spots/cells increases. Spatial transcriptomics technologies such as Visium and SMI contain thousands of spots or cells respectively, resulting in massive covariance matrices to manipulate over thousands of genes. To test for the utility of spatial models over non-spatial linear models, we randomly chose 5000 genes in each sample. Next, annotations for each ROI/spot/cell were used to indicate whether the ROI/spot/cell belonged to a biological cluster or tissue domain were. For each combination of gene and ROI/spot/cell annotation, we fit non-spatial and spatial models with exponential covariance structure to test for differential expression between the ROI/spot/cells assigned to that biological annotation and the rest ROI/spot/cells (**Table 1**). The models were fit using the *spaMM*^68^ R package on Moffitt Cancer Center’s High-Performance Computing (HPC) environment with one core assigned to each test and 8GB of memory per core. We completed AICs to compare the fit of the non-spatial and spatial models. All analyses were conducted in R (version 4.1)^66^ and visualizations with the *ggplot2*^69^ package.

## Data availability

All data sets in this study are publicly available. Please refer to Supplemental Table 1 for more information.

## Code availability

The code to conduct data pre-processing and running the models in an HPC environment can be found at https://fridleylab.github.io/diff_expression_spatial_linear_models/diff_expr_spatial_linear_models.html.

## Acknowledgments

This work has been supported in part by the National Institutes of Health (NIH) (U01 CA274489) and by the Biostatistics and Bioinformatics Shared Resource at the H. Lee Moffitt Cancer Center & Research Institute, an NCI-designated Comprehensive Cancer Center (P30 CA076292).

## Author contributions

OEO, ACS, RM, GG, XY, and BLF conceptualized and reviewed the study. BLF and OEO wrote the manuscript with contributions from all authors. OEO collected the data sets and developed the code. OEO and ACS performed the linear model tests.

## Literature Cited

1 Lein, E., Borm, L. E. & Linnarsson, S. The promise of spatial transcriptomics for neuroscience in the era of molecular cell typing. Science 358, 64–69, doi:10.1126/science.aan6827 (2017).

2 Burgess, D. J. Spatial transcriptomics coming of age. Nat Rev Genet 20, 317, doi:10.1038/s41576-019-0129-z (2019).

3 Ospina, O., Soupir, A. & Fridley, B. L. A Primer on Preprocessing, Visualization, Clustering, and Phenotyping of Barcode-Based Spatial Transcriptomics Data. Methods Mol Biol 2629, 115–140, doi:10.1007/978-1-0716-2986-4_7 (2023).

4 Chen, K. H., Boettiger, A. N., Moffitt, J. R., Wang, S. & Zhuang, X. Spatially resolved, highly multiplexed RNA profiling in single cells. Science 348, aaa6090, doi:10.1126/science.aaa6090 (2015).

5 He, S. et al. High-plex imaging of RNA and proteins at subcellular resolution in fixed tissue by spatial molecular imaging. Nat Biotechnol 40, 1794–1806, doi:10.1038/s41587-022-01483-z (2022).

6 Stahl, P. L. et al. Visualization and analysis of gene expression in tissue sections by spatial transcriptomics. Science 353, 78–82, doi:10.1126/science.aaf2403 (2016).

7 Stickels, R. R. et al. Highly sensitive spatial transcriptomics at near-cellular resolution with Slide-seqV2. Nat Biotechnol 39, 313–319, doi:10.1038/s41587-020-0739-1 (2021).

8 Cho, C. S. et al. Microscopic examination of spatial transcriptome using Seq-Scope. Cell 184, 3559–3572 e3522, doi:10.1016/j.cell.2021.05.010 (2021).

9 Moses, L. & Pachter, L. Museum of spatial transcriptomics. Nat Methods 19, 534–546, doi:10.1038/s41592-022-01409-2 (2022).

10 Seferbekova, Z., Lomakin, A., Yates, L. R. & Gerstung, M. Spatial biology of cancer evolution. Nat Rev Genet 24, 295–313, doi:10.1038/s41576-022-00553-x (2023).

11 Cheng, M. et al. Spatially resolved transcriptomics: a comprehensive review of their technological advances, applications, and challenges. J Genet Genomics 50, 625–640, doi:10.1016/j.jgg.2023.03.011 (2023).

12 Theocharidis, G. et al. Single cell transcriptomic landscape of diabetic foot ulcers. Nat Commun 13, 181, doi:10.1038/s41467-021-27801-8 (2022).

13 He, B. et al. Integrating spatial gene expression and breast tumour morphology via deep learning. Nat Biomed Eng 4, 827–834, doi:10.1038/s41551-020-0578-x (2020).

14 Stur, E. et al. Spatially resolved transcriptomics of high-grade serous ovarian carcinoma. iScience 25, 103923, doi:10.1016/j.isci.2022.103923 (2022).

15 Berglund, E. et al. Spatial maps of prostate cancer transcriptomes reveal an unexplored landscape of heterogeneity. Nat Commun 9, 2419, doi:10.1038/s41467-018-04724-5 (2018).

16 Backdahl, J. et al. Spatial mapping reveals human adipocyte subpopulations with distinct sensitivities to insulin. Cell Metab 33, 1869–1882 e1866, doi:10.1016/j.cmet.2021.07.018 (2021).

17 Andersson, A. et al. Spatial deconvolution of HER2-positive breast cancer delineates tumor-associated cell type interactions. Nat Commun 12, 6012, doi:10.1038/s41467-021-26271-2 (2021).

18 Tavares-Ferreira, D. et al. Spatial transcriptomics of dorsal root ganglia identifies molecular signatures of human nociceptors. Sci Transl Med 14, eabj8186, doi:10.1126/scitranslmed.abj8186 (2022).

19 Dhainaut, M. et al. Spatial CRISPR genomics identifies regulators of the tumor microenvironment. Cell 185, 1223–1239 e1220, doi:10.1016/j.cell.2022.02.015 (2022).

20 Garcia-Alonso, L. et al. Mapping the temporal and spatial dynamics of the human endometrium in vivo and in vitro. Nat Genet 53, 1698–1711, doi:10.1038/s41588-021-00972-2 (2021).

21 Chen, H. et al. Dissecting mammalian spermatogenesis using spatial transcriptomics. Cell Rep 37, 109915, doi:10.1016/j.celrep.2021.109915 (2021).

22 Delorey, T. M. et al. COVID-19 tissue atlases reveal SARS-CoV-2 pathology and cellular targets. Nature 595, 107–113, doi:10.1038/s41586-021-03570-8 (2021).

23 Rao, A., Barkley, D., Franca, G. S. & Yanai, I. Exploring tissue architecture using spatial transcriptomics. Nature 596, 211–220, doi:10.1038/s41586-021-03634-9 (2021).

24 Longo, S. K., Guo, M. G., Ji, A. L. & Khavari, P. A. Integrating single-cell and spatial transcriptomics to elucidate intercellular tissue dynamics. Nat Rev Genet 22, 627–644, doi:10.1038/s41576-021-00370-8 (2021).

25 Marshall, J. L. et al. High Resolution Slide-seqV2 Spatial Transcriptomics Enables Discovery of Disease-Specific Cell Neighborhoods and Pathways. iScience, 104097, doi:10.1016/j.isci.2022.104097 (2022).

26 Joshi, N. et al. A spatially restricted fibrotic niche in pulmonary fibrosis is sustained by M-CSF/M-CSFR signalling in monocyte-derived alveolar macrophages. Eur Respir J 55, doi:10.1183/13993003.00646-2019 (2020).

27 Su, S. & Li, X. Dive into Single, Seek Out Multiple: Probing Cancer Metastases via Single-Cell Sequencing and Imaging Techniques. Cancers (Basel) 13, doi:10.3390/cancers13051067 (2021).

28 Fang, S. et al. Computational Approaches and Challenges in Spatial Transcriptomics. Genomics Proteomics Bioinformatics, doi:10.1016/j.gpb.2022.10.001 (2022).

29 Ren, Y. et al. Spatial transcriptomics reveals niche-specific enrichment and vulnerabilities of radial glial stem-like cells in malignant gliomas. Nat Commun 14, 1028, doi:10.1038/s41467-023-36707-6 (2023).

30 Lyubetskaya, A. et al. Assessment of spatial transcriptomics for oncology discovery. Cell Rep Methods 2, 100340, doi:10.1016/j.crmeth.2022.100340 (2022).

31 Zhu, J. et al. Delineating the dynamic evolution from preneoplasia to invasive lung adenocarcinoma by integrating single-cell RNA sequencing and spatial transcriptomics. Exp Mol Med 54, 2060–2076, doi:10.1038/s12276-022-00896-9 (2022).

32 Buzzi, R. M. et al. Spatial transcriptome analysis defines heme as a hemopexin-targetable inflammatoxin in the brain. Free Radic Biol Med 179, 277–287, doi:10.1016/j.freeradbiomed.2021.11.011 (2022).

33 Luo, W. et al. Single-cell spatial transcriptomic analysis reveals common and divergent features of developing postnatal granule cerebellar cells and medulloblastoma. BMC Biol 19, 135, doi:10.1186/s12915-021-01071-8 (2021).

34 Qiu, Z. et al. Detection of differentially expressed genes in spatial transcriptomics data by spatial analysis of spatial transcriptomics: A novel method based on spatial statistics. Front Neurosci 16, 1086168, doi:10.3389/fnins.2022.1086168 (2022).

35 Svensson, V., Teichmann, S. A. & Stegle, O. SpatialDE: identification of spatially variable genes. Nat Methods 15, 343–346, doi:10.1038/nmeth.4636 (2018).

36 Fornito, A., Arnatkeviciute, A. & Fulcher, B. D. Bridging the Gap between Connectome and Transcriptome. Trends Cogn Sci 23, 34–50, doi:10.1016/j.tics.2018.10.005 (2019).

37 Su, J. et al. Smoother: a unified and modular framework for incorporating structural dependency in spatial omics data. Genome Biol 24, 291, doi:10.1186/s13059-023-03138-x (2023).

38 Edsgard, D., Johnsson, P. & Sandberg, R. Identification of spatial expression trends in single-cell gene expression data. Nat Methods 15, 339–342, doi:10.1038/nmeth.4634 (2018).

39 Hu, J. et al. SpaGCN: Integrating gene expression, spatial location and histology to identify spatial domains and spatially variable genes by graph convolutional network. Nat Methods 18, 1342–1351, doi:10.1038/s41592-021-01255-8 (2021).

40 Zhu, J., Sun, S. & Zhou, X. SPARK-X: non-parametric modeling enables scalable and robust detection of spatial expression patterns for large spatial transcriptomic studies. Genome Biol 22, 184, doi:10.1186/s13059-021-02404-0 (2021).

41 Miller, B. F., Bambah-Mukku, D., Dulac, C., Zhuang, X. & Fan, J. Characterizing spatial gene expression heterogeneity in spatially resolved single-cell transcriptomic data with nonuniform cellular densities. Genome Res 31, 1843–1855, doi:10.1101/gr.271288.120 (2021).

42 Dries, R. et al. Giotto: a toolbox for integrative analysis and visualization of spatial expression data. Genome Biol 22, 78, doi:10.1186/s13059-021-02286-2 (2021).

43 Weber, L. M., Saha, A., Datta, A., Hansen, K. D. & Hicks, S. C. nnSVG for the scalable identification of spatially variable genes using nearest-neighbor Gaussian processes. Nat Commun 14, 4059, doi:10.1038/s41467-023-39748-z (2023).

44 Deshpande, A. et al. Uncovering the spatial landscape of molecular interactions within the tumor microenvironment through latent spaces. Cell Syst 14, 285–301 e284, doi:10.1016/j.cels.2023.03.004 (2023).

45 Chen, C., Kim, H. J. & Yang, P. Evaluating spatially variable gene detection methods for spatial transcriptomics data. Genome Biol 25, 18, doi:10.1186/s13059-023-03145-y (2024).

46 Squair, J. W. et al. Confronting false discoveries in single-cell differential expression. Nat Commun 12, 5692, doi:10.1038/s41467-021-25960-2 (2021).

47 Park, Y. P. & Kellis, M. CoCoA-diff: counterfactual inference for single-cell gene expression analysis. Genome Biol 22, 228, doi:10.1186/s13059-021-02438-4 (2021).

48 Soneson, C. & Robinson, M. D. Bias, robustness and scalability in single-cell differential expression analysis. Nat Methods 15, 255–261, doi:10.1038/nmeth.4612 (2018).

49 Robinson, M. D., McCarthy, D. J. & Smyth, G. K. edgeR: a Bioconductor package for differential expression analysis of digital gene expression data. Bioinformatics 26, 139–140, doi:10.1093/bioinformatics/btp616 (2010).

50 Love, M. I., Huber, W. & Anders, S. Moderated estimation of fold change and dispersion for RNA-seq data with DESeq2. Genome Biol 15, 550, doi:10.1186/s13059-014-0550-8 (2014).

51 Cressie, N. A. C. Statistics for spatial data. 900 (Wiley & Sons, 1993).

52 Pinheiro, J. C. & Bates, D. M. Mixed-Effects Models in S and S-PLUS. (Springer, 2000).

53 Hafemeister, C. & Satija, R. Normalization and variance stabilization of single-cell RNA-seq data using regularized negative binomial regression. Genome Biol 20, 296, doi:10.1186/s13059-019-1874-1 (2019).

54 Lun, A. T., McCarthy, D. J. & Marioni, J. C. A step-by-step workflow for low-level analysis of single-cell RNA-seq data with Bioconductor. F1000Res 5, 2122, doi:10.12688/f1000research.9501.2 (2016).

55 Bergholtz, H. et al. Best Practices for Spatial Profiling for Breast Cancer Research with the GeoMx((R)) Digital Spatial Profiler. Cancers (Basel) 13, doi:10.3390/cancers13174456 (2021).

56 Zhao, P., Zhu, J., Ma, Y. & Zhou, X. Modeling zero inflation is not necessary for spatial transcriptomics. Genome Biol 23, 118, doi:10.1186/s13059-022-02684-0 (2022).

57 Jiang, X., Xiao, G. & Li, Q. A Bayesian modified Ising model for identifying spatially variable genes from spatial transcriptomics data. Stat Med 41, 4647–4665, doi:10.1002/sim.9530 (2022).

58 Li, Q., Zhang, M., Xie, Y. & Xiao, G. Bayesian Modeling of Spatial Molecular Profiling Data via Gaussian Process. Bioinformatics, doi:10.1093/bioinformatics/btab455 (2021).

59 Fast gene set enrichment analysis v. 1.26 (Bioconductor, 2019).

60 Liberzon, A. et al. The Molecular Signatures Database (MSigDB) hallmark gene set collection. Cell Syst 1, 417–425, doi:10.1016/j.cels.2015.12.004 (2015).

61 Ravi, V. M. et al. Spatially resolved multi-omics deciphers bidirectional tumor-host interdependence in glioblastoma. Cancer Cell 40, 639–655 e613, doi:10.1016/j.ccell.2022.05.009 (2022).

62 Zhu, G., Pei, L., Xia, H., Tang, Q. & Bi, F. Role of oncogenic KRAS in the prognosis, diagnosis and treatment of colorectal cancer. Mol Cancer 20, 143, doi:10.1186/s12943-021-01441-4 (2021).

63 spatialGE: An R package for visualization and analysis of spatially-resolved gene expression v. 1.2 (GitHub, 2023).

64 Maynard, K. R. et al. Transcriptome-scale spatial gene expression in the human dorsolateral prefrontal cortex. Nat Neurosci 24, 425–436, doi:10.1038/s41593-020-00787-0 (2021).

65 Nanostring.

66 R: A language and environment for statistical computing v. v4.1.2 (R Foundation for Statistical Computing, Viena, Austria, 2021).

67 Ospina, O. E. et al. spatialGE: Quantification and visualization of the tumor microenvironment heterogeneity using spatial transcriptomics. Bioinformatics 38, 2645–2647, doi:10.1093/bioinformatics/btac145 (2022).

68 Rousset, F. & Ferdy, J.-B. Testing environmental and genetic effects in the presence of spatial autocorrelation. Ecography 37, 781–790, doi:10.1111/ecog.00566 (2014).

69 Wickham, H. ggplot2: Elegant Graphics for Data Analysis. (Springer-Verlag New York, 2016).

